# Induction of RNA interference by *C. elegans* mitochondrial dysfunction via the DRH-1/RIG-I homologue RNA helicase and the EOL-1/RNA decapping enzyme

**DOI:** 10.1101/2020.06.05.136978

**Authors:** Kai Mao, Peter Breen, Gary Ruvkun

**Affiliations:** Department of Molecular Biology, Massachusetts General Hospital, Boston, MA 02114, USA; Department of Genetics, Harvard Medical School, Boston, MA 02115, USA

## Abstract

RNA interference (RNAi) is an antiviral pathway common to many eukaryotes that detects and cleaves foreign nucleic acids. In mammals, mitochondrially localized proteins such as MAVS, RIG-I, and MDA5 mediate antiviral responses. Here, we report that mitochondrial dysfunction in *Caenorhabditis elegans* activates RNAi-directed silencing via induction of a pathway homologous to the mammalian RIG-I helicase viral response pathway. The induction of RNAi also requires the conserved RNA decapping enzyme EOL-1/DXO. The transcriptional induction of *eol-1* requires DRH-1 as well as the mitochondrial unfolded protein response (UPR^mt^). Upon mitochondrial dysfunction, EOL-1 is concentrated into foci that depend on the transcription of mitochondrial RNAs that may form dsRNA, as has been observed in mammalian antiviral responses. The enhanced RNAi triggered by mitochondrial dysfunction contributes to the increase in longevity that is induced by mitochondrial dysfunction.

## Introduction

Many RNA viruses carry an RNA-dependent RNA polymerase (RdRp) to replicate their RNA genome in the host, bypassing entirely information storage and replication with DNA. In this way, RNA viruses can replicate in non-dividing cells with lower deoxyribonucleotide levels than those required for DNA viruses or retroviruses but with the substantial ribonucleotide levels needed for transcription of RNA and mRNAs in terminally differentiated cells. A double-stranded RNA (dsRNA) viral replication intermediate is a strong clue to the host cell that an RNA virus infection is underway. Recognition of the dsRNA replication intermediate is an initial step in antiviral immune responses. In mammals, the RNA helicases RIG-I or MDA5 recognize the RNA signatures of RNA virus replication (Chow et al., 2018). RIG-I binds to dsRNA, as well as single-stranded RNA (ssRNA) with 5’-triphosphate that is a signature of the products from RNA dependent RNA polymerases that mediate viral RNA replication (Lassig and Hopfner, 2017). RIG-I or MDA5 associate with the mitochondrial antiviral signaling protein (MAVS) and elicit the downstream NFkappaB and other interferon immune signaling that mediate systemic antiviral immune defenses (Zevini et al., 2017).

The nematode *Caenorhabditis elegans* uses an RNA interference (RNAi) pathway to mediate antiviral defense instead of the interferon signaling of vertebrates and most invertebrates (Felix et al., 2011; Lu et al., 2005; Schott et al., 2005; Wilkins et al., 2005). RNA interference is a highly conserved mechanism for antiviral defense and broadly deployed by eukaryotes, including, fungi, nematodes, insects, plants and vertebrates (Ding et al., 2018; Gammon and Mello, 2015; Szittya and Burgyan, 2013). RNA interference was initially discovered to mediate silencing triggered by engineered double-stranded RNA triggers but soon discovered to produce natural small interfering RNAs (siRNAs) that target specific mRNAs for degradation. RNA interference is mediated by 22 nt to 26 nt single stranded short interfering or siRNAs that are produced and presented to target mRNAs by the Argonaute proteins and the Dicer dsRNA ribonuclease, conserved across many but not all eukaryotes (Tabach et al., 2013). Many species of fungi and plants and a few animal species such as nematodes, ticks, and scorpions, have a second stage amplifier for siRNAs, RNA dependent RNA polymerases that extend off of the primary siRNAs produced by Dicer and Argonaute proteins to render their RNAi pathways especially potent. In fact, RNAi was simultaneously discovered in fungal, plant, and nematode strains that all possess such RdRp genes to potentiate their RNAi response pathways. RNA viruses of course depend on such RdRp genes for their replication so the defense pathway may have evolved from the viral weapon, or vice versa. The fact that RdRp genes mediate RNAi functions across eukaryotic phylogeny suggests that these pathways have been lost for example in most of the vertebrate lineages that only have the first stage of RNAi, the Argonautes and Dicer, rather than RdRps evolving or being acquired by horizontal transfer from viruses independently in so many eukaryotic lineages. The Sting and NFkappaB RNA virus surveillance pathway of vertebrates and many invertebrates are anti-correlated with the presence of RdRp genes in the genomes of the peculiar animals that have retained them and may have displaced the RdRp second stage of the RNAi pathway.

Some of the genes in the vertebrate and insect NFkappaB pathway are conserved between nematodes and other animals without RdRp genes. Like its mammalian orthologues RIG-I and MDA5, the *C. elegans* DRH-1 mediates antiviral RNAi and is essential for neutralization of an invading virus (Gammon et al., 2017; Guo et al., 2013). DRH-1 was initially identified as a binding partner with the Dicer protein DCR-1, the Argonaute protein RDE-1 and the RNA helicase RDE-4 (Tabara et al., 2002). In addition to *drh-1, drh-2* is a pseudogene, and *drh-3* encodes a component of an RdRP protein complex (Duchaine et al., 2006).

The DCR-1 ribonuclease is the first step in siRNA generation in most of the *C. elegans* RNAi pathways, including exogenous RNAi, endogenous RNAi and antiviral RNAi (Ashe et al., 2013; Duchaine et al., 2006). siRNAs generated by these pathways engage the Mutator proteins that mediate siRNA-guided repression in both exogenous and endogenous RNAi (Phillips et al., 2012; Zhang et al., 2011). Mutations in the Dicer, Argonaute, and accessory factors that generate and present siRNAs to target mRNAs are generally defective for RNAi or antiviral defense. But the Argonaute and RdRp gene families are expanded in *C. elegans* compared to most animals, and surprisingly, loss of function mutations in some of those genes cause an increase in the response to siRNAs: mutations in the RdRp RRF-3, the specialized Argonaute ERGO-1, the RNA helicase ERI-6/7, or the exoribonuclease ERI-1 enhance silencing by siRNAs (Fischer et al., 2008; Kennedy et al., 2004). Many of these enhanced RNAi mutations desilence retroviral elements in the *C. elegans* genome so that antiviral response pathways are triggered to in turn induce the expression of RNAi-based antiviral defense (Fischer and Ruvkun, 2020).

A mutation in the *C. elegans* mitochondrial chaperone gene *hsp-6* causes induction of a suite of drug detoxification and defense genes (Mao et al., 2019). Here we show that in addition to the induction of these defense pathways, RNA interference pathways are also activated. Because RNA interference is a key feature of *C. elegans* antiviral defense, and because of the association of mammalian viral defense pathways such as MAVS and RIG-I with the mitochondrion, we explored more fully how mitochondrial homeostatic pathways connect to RNA interference and antiviral defense in *C. elegans*. We find that reduction of function mutations in a wide range of mitochondrial components robustly enhanced RNA interference-mediated silencing of endogenous genes as well as a variety of reporters of RNAi. These antiviral responses to mitochondrial dysfunction are homologous to the RIG-I-based mitochondrial response in mammals because they depend on the RIG-I homologue, the DRH-1 RNA helicase. Comparing the *C. elegans* transcriptional response of a mitochondrial mutant and infection with the Orsay RNA virus, we found a striking overlap of expression of multiple members of *C. elegans pals-* genes implicated in anti-viral and anti-pathogen response pathways (Leyva-Diaz et al., 2017; Reddy et al., 2017) and the *eol-1*/DXO RNA decapping enzyme gene.

We found that *eol-1* transcription is dramatically induced by mitochondrial dysfunction, and an *eol-1* null mutation in strongly suppresses the DRH-1-mediated antiviral RNAi response normally induced by mitochondrial dysfunction. We showed that the EOL-1 protein forms foci in the cytosol only if the mitochondrion is stressed, and the production of these foci are dependent on production of RNA from the mitochondrial genome. This is reminiscent of the central role that dsRNA released from the mitochondria plays in mammalian antiviral response pathways. During a viral infection, mitochondrial disruption by MAVS and other mitochondrial-associated viral immunity factors releases mitochondrial dsRNA and DNA to the cytosol, where dsRNAs trigger an MDA5-dependent interferon response (Dhir et al., 2018; Pajak et al., 2019) and DNA via cGAS and Sting activate NFkappaB immune signaling (West et al., 2015).

Gene inactivation or mutations in a wide variety of nuclear-encoded mitochondrial genes causes one of the strongest increases in lifespan in genome screens for lifespan extension (Lee et al., 2003). We find that the uncoupling the antiviral defense response via *drh-1* or *eol-1* mutations abrogates the increase in lifespan, suggesting that the antiviral axis of mitochondrial dysfunction is critical to the lifespan extension.

## Results

### *C. elegans* mitochondrial mutations enhance RNA interference using the RNA helicase DRH-1

The *hsp-6(mg585)* allele is a reduction of function mutation (P386S) in the mitochondrial HSP70 chaperone in a region that is conserved between mammals and *C. elegans* (VQEIFGKV**P**SKAVNPDEAVA). *hsp-6(mg585)* causes induction of a suite of drug detoxification and defense genes that are also induced by a variety of mitochondrial mutations or toxins that disrupt mitochondrial function (Mao et al., 2019). *C. elegans* HSP-6 and human mtHSP70 are orthologues of bacterial and archaeal dnaK; these mitochondrial chaperones were bacterial chaperones before the mitochondrial endosymbiosis event more than a billion years ago and the migration of these bacterial genes to the eukaryotic nuclear genome (Mao et al., 2019). Null alleles of *hsp-6* cause developmental arrest, presumably because of the defects in the folding or import of many mitochondrial client proteins (Kim et al., 2016; Wiedemann and Pfanner, 2017). But the viable *hsp-6(mg585)* allele allows mitochondrial roles in other pathways to be studied without the associated pleiotropic lethality. In addition to activating detoxification and immune responses, the mitochondrial defects caused by *hsp-6(mg585)* also unexpectedly cause enhanced RNA interference which is an antiviral defense pathway.

RNA interference in *C. elegans* is gene silencing by mRNA degradation or heterochromatin induction that can be induced by injection or ingestion of approximately 1 Kb of double stranded RNA corresponding to a particular gene. For feeding RNAi, *E. coli* have been engineered to produce any of 18,000 dsRNAs corresponding to any *C. elegans* gene (Kim et al., 2005). The specificity of RNAi in *C. elegans* is superior to the single siRNA approaches common in most animal systems, probably because, almost unique to animals, *C. elegans* uses RNA-dependent RNA polymerase genes to amplify the primary short interfering RNAs (siRNAs) produced by the Argonaute and Dicer proteins that nearly all eukaryotes also use for RNAi, and unlike in other animal systems, RNA interference in *C. elegans* can be induced by 1 Kb segments of dsRNA without the induction of interferon-related responses to dsRNA. Thus thousands of siRNAs produced from feeding *C. elegans* with an *E. coli* expressing a 1 Kb dsRNA sum for on-target effects on mRNA inactivation and average for off-target mRNA inactivations. The specificity of dsRNA-induced RNAi was validated with genetic loci that had previously studied by genetics: most dsRNAs corresponding to those genes with known phenotypes recapitulated the phenotypes predicted from genetics. But a minority of genes expected to generate easily scored phenotypes by RNA interference did not silence in wild type. These dsRNAs were used in genetic screens for *C. elegans* mutations that enhance RNAi, or eri-mutations (Kennedy et al., 2004). For example, mutations in the conserved exonuclease *eri-1* or the RdRp *rrf-3* cause enhanced RNAi (Duchaine et al., 2006). Most mutants that enhance RNAi responses also cause silencing of transgenes, because the transgene is detected as a bearing some foreign signatures (lack of introns is a major signature of a viral origin) and silenced in the enhanced RNAi state of these mutant strains. A set of dsRNA tester genes have been developed that cause strong phenotypes in enhanced RNAi mutants, but no RNAi phenotype in wild type (Guang et al., 2008).

We found that the mitochondrial *hsp-6(mg585)* mutant showed the strong phenotype seen in enhanced RNAi mutants on many of these enhanced RNAi tester dsRNAs expressed from *E. coli*. For example, a dsRNA targeting the *lir-1* (where *lir* is an abbreviation for *lin-26*-related) gene causes no phenotype in wild type, but causes lethal arrest in for example the *eri-6(mg379)* enhanced RNAi mutant, as well as on the mitochondrial *hsp-6(mg585)* mutant (Figure 1A). The enhanced lethality after exposure to *lir-1* dsRNA in enhanced RNAi mutants is due to siRNAs from this dsRNA targeting the duplicated and diverged genes with high regions of nucleotide homology on the same primary transcript of the *lir-1, lir-2*, and *lin-26* operon (Pavelec et al., 2009). This enhanced response to *lir-1* was not due to strain background mutations in *hsp-6(mg585)* or the pleiotropy of a mitochondrial chaperone that may affect the function of many imported mitochondrial proteins: *lir-1* RNAi also caused a lethal arrest on a variety of other nuclearly-encoded mitochondrial protein point mutants, including NADH dehydrogenase *nuo-6*/NDUFB4, ubiquinone biosynthesis *clk-1*/COQ7, and the iron-sulfur cluster *isp-1*/UQCRFS1 (Figure 1A). Many mutations in nuclearly-encoded mitochondrial proteins are lethal, so that amino acid substitution reduction in function alleles constitute most of the available viable mitochondrial mutations. A distinct dsRNA that targets a histone 2B gene, *his-44*, causes larval arrest in enhanced RNAi mutants but no lethality on wild type (Wang and Ruvkun, 2004) also caused larval arrest in the *hsp-6/mtHSP70, nuo-6*/NDUFB4, *clk-1*/COQ7, and *isp-1*/UQCRFS1 mitochondrial mutants (Figure 1A). *his-44* maps to a cluster of histone genes including multiple histone 2B genes with nucleic acid homology; in the enhanced RNAi mutants, the initial siRNAs produced from the *his-44* dsRNA may spread to adjacent histone 2B genes.

**Figure 1.**
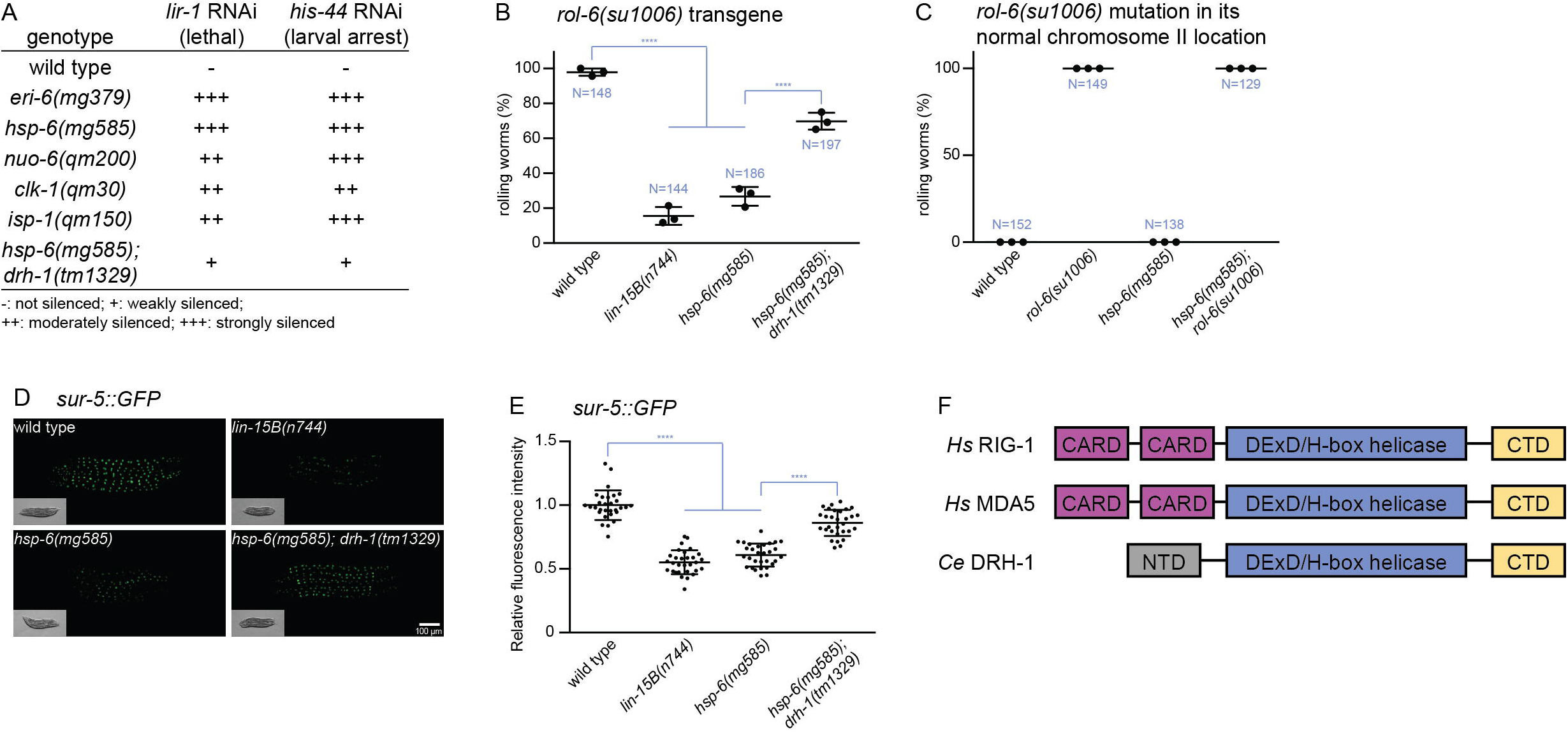
Mitochondrial mutants show enhanced RNAi through DRH-1. (A) Enhanced RNAi response to *lir-1* RNAi or *his-44* RNAi in the *eri-6* control enhanced RNAi mutant or any of the mitochondrial mutants (*hsp-6, nuo-6, clk-1, or isp-1*), causes lethality/arrest on the mitochondrial mutants but not wild type. The enhanced RNAi of the *hsp-6* mitochondrial mutant is suppressed by a *drh-1* mutation. (B) Transgene silencing test with *rol-6(su1006)* transgene. This transgene causes a Rolling behavior due to a twist on the collagen cuticle in wild type. The expression of the *rol-6* collagen mutation from the transgene is silenced by enhanced RNAi in mitochondrial mutants, and this transgene silencing depends on *drh-1* gene activity. Results of 3 replicate experiments are shown. N: total number; ****p<0.0001. (C) The mitochondrial mutations do not simply suppress the collagen defect of a *rol-6* mutation: if the *rol-6(su1006)* mutation is not on a transgene that is under RNAi control, the Rol phenotype is not suppressed by *hsp-6(mg585)* and an *hsp-6(mg585)* single mutation does not have any Rol phenotype. Results of 3 replicate experiments are shown. N: total number. (D) and (E) Transgene silencing test with *sur-5::GFP* transgene. This transgene is ubiquitously expressed in all somatic cells. The expression of the *sur-5::GFP* from the transgene is silenced by enhanced RNAi in mitochondrial mutants, and this transgene silencing depends on *drh-1* gene activity. Animals were imaged in (D) and the fluorescence was quantified in (E). ****p<0.0001. (F) Diagram of human RIG-I, MDA5 and *C. elegans* DRH-1. The helicase domain and CTD are conserved. CARD: caspase recruitment domain; NTD: N-terminal domain; CTD: C-terminal domain; *Hs*: Homo sapiens; *Ce*: *Caenorhabditis elegans*.

One explanation for the enhanced response to *lir-1* and *his-44* RNAi is that these genes have genetic interactions with *hsp-6(mg585)* that caused synthetic enhancement of certain phenotypes. However because multiple mitochondrial mutations enhance response to these enhanced RNAi tester dsRNAs, another hypothesis was that *hsp-6(mg585)* and the other mitochondrial mutants trigger an enhanced RNAi (Eri) phenotype. Three other mitochondrial point mutants that we tested (out of six tested), *nduf-7(et19), mev-1(kn1)* and *gas-1(fc21)* did not cause an enhanced RNAi phenotype. There was no obvious distinction between the types of mitochondrial mutations that caused enhanced RNAi and those that did not, in terms of growth rate or severity of the phenotype. NUO-6, GAS-1, and NDUF-7 are components of complex I, MEV-1 is from complex II, ISP-1 is a component of complex III, and CLK-1 produces the ubiquinone that also functions in electron transport. It is possible that only particular mitochondrial insults activate the RNAi pathway. But because this was not a rare response to a peculiar mitochondrial dysfunction, but rather associated with many different mitochondrial mutations we explored this with other measures of the intensity of RNA interference.

A hallmark of an enhanced RNAi phenotype is the silencing of transgenes, as increased RNA interference detects the foreign genetic signatures (the fusion of non-*C. elegans* genes such as GFP, and the synthetic introns, and other engineered features) of transgenes (Fischer et al., 2013). We asked if the *hsp-6(mg585)* mutant displayed enhanced transgene silencing by introducing a transgene containing *rol-6(su1006)*, a dominant mutation of a hypodermal collagen that causes an easily scored rolling movement phenotype (Kramer and Johnson, 1993). *lin-15B* is a class B synthetic multivulva (synMuv B) gene and loss-of-function of *lin-15B* significantly enhances transgene silencing (Wang et al., 2005). A chromosomally-integrated transgene carrying multiple copies of the *rol-6(su1006)* mutant collagen gene causes 100% of transgenic animals to roll (a Rol phenotype) in wild type animals, but in strains with enhanced RNA interference, this transgene is now silenced so that 16% of animals are Rol in the *lin-15B(n744)* mutant and 27% Rol in the *hsp-6(mg585)* homozygous mutant (Figure 1B). The suppression of the Rol phenotype from the transgene carrying *rol-6(su1006)* was not due to a genetic interaction between *hsp-6(mg585)* and the *rol-6(su1006)* mutant collagen gene, because the *hsp-6(mg585)*; *rol-6(su1006)* double mutant with the *rol-6* collagen mutation located in its normal chromosomal location not subject to enhanced RNAi silencing of a transgene, was still 100% Rol (Figure 1C). Rather, the enhanced RNAi of the mitochondrial mutants, including *hsp-6(mg585)* causes a silencing of the *rol-6(su1006)* mutant collagen allele on the multicopy transgene to suppress the Rol phenotype.

To evaluate transgene silencing in other tissues, a ubiquitously expressed *sur-5::GFP* fusion gene was monitored in all somatic cells (Yochem et al., 1998). The bright GFP signal of *sur-5::GFP* was dramatically decreased in the enhanced RNAi mutant *lin-15B(n744)* as well as in the *hsp-6(mg585)* mitochondrial mutant (Figure 1D and 1E). Thus *hsp-6(mg585)* enhances exogenous RNAi and silences somatic transgenes. The somatic transgene silencing and enhanced RNAi in the *eri-1* mutant is associated with a failure to nuclearly-localize the Argonaute trancriptional silencing factor NRDE-3 that acts downstream of siRNA generation (Guang et al., 2008; Wu et al., 2012). The synMuvB enhanced RNAi mutants cause a somatic misexpression of the normally germline P granules implicated in siRNA (Wu et al., 2012). However, neither NRDE-3 nuclear delocalization nor the somatic expression of P granules occurred in *hsp-6(mg585)* mutant, suggesting that mitochondrial dysfunction does not induce *eri-1* or synMuvB classes of enhanced RNAi.

The enhanced RNAi of *hsp-6(mg585)* most resembled the *eri-6/7* RNA helicase and *ergo-1* Argonaute enhanced RNAi phenotypes, associated with desilencing of recently acquired viral genes and induction of viral immunity, without nuclear NRDE-3 or somatic P granule expression (Fischer et al., 2011; Fischer et al., 2013; Fischer and Ruvkun, 2020). In mammalian cells, the RNA helicase MDA5 mediates an interferon antiviral immune response that is strongly enhanced by a mitochondrial RNA degradation mutation that enhances production of mitochondrial dsRNAs (Dhir et al., 2018). Intriguingly, mutations in the *C. elegans* homologue of MDA5, *drh-1* suppress the synthetic lethality of enhanced RNAi and dsRNA editing double mutants (Fischer and Ruvkun, 2020; Reich et al., 2018). *C. elegans* DRH-1 contains three domains, including conserved helicase domain and C-terminal domain (CTD) (Figure 1F), and an N-terminal domain (NTD) that is only conserved in nematodes (Supplemental Figure 1). The caspase activation and recruitment domains (CARD) of RIG-I and MDA5 associate with the mitochondrial protein MAVS, which is unique to mammals (Zevini et al., 2017). Conversely, the NTD of DRH-1 is nematode-specific and essential for the inhibition of viral replication (Guo et al., 2013). To test if DRH-1 is required for *hsp-6(mg585)* induced transgene silencing, two transgenes *rol-6(su1006)* and *sur-5::GFP* were tested for silencing in an *hsp-6; drh-1* double mutant. The *drh-1(tm1329)* allele disrupts the NTD and renders the strain susceptible to viral infection (Gammon et al., 2017). *drh-1(tm1329)* suppressed the transgene silencing induced by *hsp-6(mg585)*: *hsp-6(mg585)* animals carrying the *rol-6(su1006)* transgene were 27% Rol, but the *hsp-6(mg585); drh-1(tm1329)* double mutant was 70% Rol (Figure 1B), showing that the transgene silencing induced by *hsp-6(mg585)* was suppressed by *drh-1(tm1329)*. Similarly, the GFP signal of *sur-5::GFP* was significantly brighter in the *hsp-6(mg585); drh-1(tm1329)* double mutant compared to *hsp-6(mg585)* (Figure 1D and 1E). The *lir-1* and *his-44* enhanced RNAi lethal or larval arrest phenotypes of *hsp-6(mg585)* were also suppressed by *drh-1(tm1329)* (Figure 1A). *drh-1* mutant animals carrying a distinct mutant allele from a wild strain of *C. elegans* with a C-terminal truncation are competent for RNAi but show defects in antiviral RNAi (Ashe et al., 2013). Therefore, *hsp-6(mg585*) enhanced silencing of transgenes and of *his-44* and *lir-1* enhanced RNAi requires DRH-1 gene activity.

### EOL-1 acts downstream of DRH-1 for somatic silencing

Upon virus infection, human RIG-I and MDA5 mediate the upregulation of interferon genes. Although the interferon signaling pathway is not conserved in *C. elegans*, we suspected DRH-1, like its mammalian orthologue, might promote the transcriptional activation of downstream response genes. Comparison of the expression profile of *hsp-6(mg585)* (Mao et al., 2019) and wild type animals infected with the RNA virus Orsay (Chen et al., 2017) showed that of the 126 genes upregulated by Orsay virus infection, 45 were also upregulated in *hsp-6(mg585)* mutant (Figure 2A and Supplemental Table S1). Of these 45 genes, 12 are members of the *pals-1* to *pals-40* genes, which located in a few clusters and, remarkably, a null allele in *pals-22* causes an enhanced RNAi phenotype of transgene silencing. Thus the increased expression of *pals-* genes in virus infected and mitochondrial defective animals is not just associated with enhanced RNA interference but a mutation in one *pals-22* gene can cause increased RNA interference and transgene silencing (Leyva-Diaz et al., 2017; Reddy et al., 2017).

**Figure 2.**
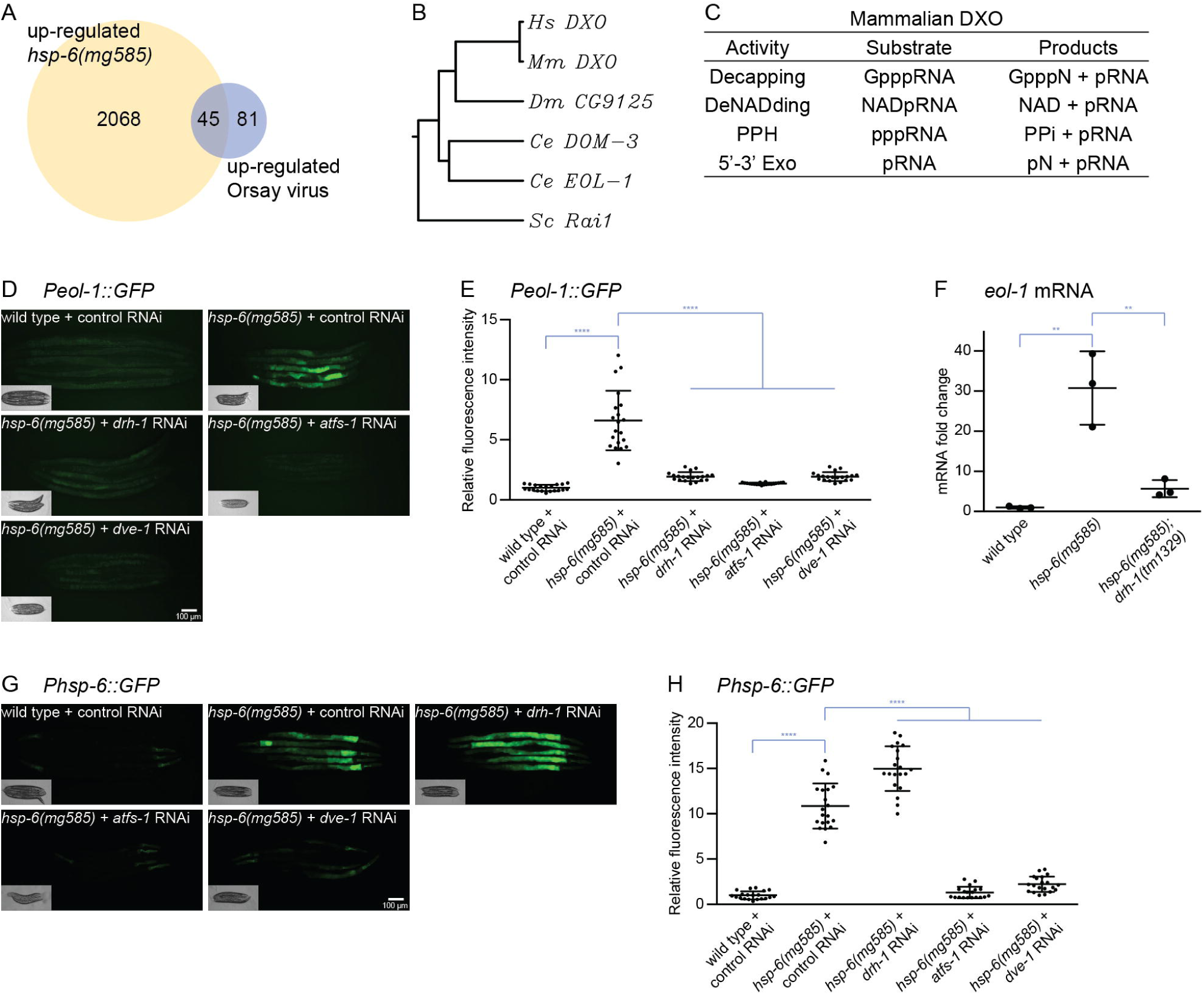
Increased *eol-1* expression depends on DRH-1. (A) Venn diagram for genes upregulated in *hsp-6(mg585)* and Orsay virus infection. (B) Phylogenetic tree of EOL-1/DXO. EOL-1 is conserved from yeast to mammals. *Hs*: Homo sapiens; *Mm*: *Mus musculus*; *Dm*: *Drosophila melanogaster*; *Ce*: *Caenorhabditis elegans*; *Sc*: *Saccharomyces cerevisiae*. (C) The enzyme activities of human DXO. Mammalian DXO modifies the 5’ end of mRNAs: decapping, deNADing, pyrophosphohydrolase and 5’-3’ exonuclease. (D) and (E) The induction of *Peol-1::GFP* transcriptional fusion reporter requires *drh-1* and UPR^mt^. The *Peol-1::GFP* is strongly induced by *hsp-6(mg585)* mitochondrial mutant, and this induction is abrogated by *drh-1* RNAi as well as RNAi of genes involved in UPR^mt^ (*atfs-1* and *dve-1*). Animals were imaged in (D) and the fluorescence was quantified in (E). ****p<0.0001. (F) The *hsp-6(mg585)* mutant causes increased mRNA level of *eol-1* in a *drh-1* dependent manner. RT-qPCR assays showed the mRNA level of *eol-1* was induced by *hsp-6(mg585)* mutant and this induction was abolished in *hsp-6(mg585); drh-1(tm1329)* double mutant. **p<0.01. (G) and (H) DRH-1 does not contribute to UPR^mt^. The benchmark reporter of UPR^mt^ *Phsp-6::GFP* is induced by the *hsp-6(mg585)* mutant. And the induction is suppressed by RNAi of *atfs-1* or *dev-1*, but not *drh-1*. Animals were imaged in (G) and the fluorescence was quantified in (H). ****p<0.0001.

The *eol-1* RNA decapping gene was also strongly induced in virally infected and mitochondrial mutant *C. elegans* (Supplemental Table S1). We selected *eol-1* for detailed analysis because a. it is conserved from yeast to mammals (Figure 2B), b. *eol-1* expression is induced by multiple mitochondrial mutations as well as in a *eri-6; adr-1/2* enhanced RNAi mutant that desilences endogenous retroviruses and retrotransposons (Fischer and Ruvkun, 2020; Senchuk et al., 2018), c. the induction of *eol-1* by Orsay virus infection is dependent on *drh-1* (Sowa et al., 2020), and d. the human orthologue of EOL-1, DXO represses hepatitis C virus replication (Amador-Canizares et al., 2018). *C. elegans eol-1* was initially identified as a mutant that enhanced olfactory learning after *Pseudomonas aeruginosa* infection (Shen et al., 2014). *eol-1* encodes a decapping exoribonulease orthologous to yeast Rai1 and mammalian DXO (Figure 2B). Mammalian DXO processes the 5’ end of mRNA (Kramer and McLennan, 2019), including a decapping activity that removes the unmethylated guanosine cap and the first nucleotide (GpppN), the deNADing activity that removes the nicotinamide adenine dinucleotide (NAD) cap, the pyrophosphohydrolase (PPH) activity that releases pyrophosphate (PPi) from 5’ triphosphorylated RNA, and 5’-3’ exonuclease (5’-3’ Exo) to degrade the entire RNA (Figure 2C). The removal the m^7^G cap and subsequent degradation of mammalian mRNAs are directed by the decapping exoribonuclease DXO. Expression of *Mus musculus* DXO rescued the enhanced olfactory learning phenotype in *C. elegans eol-1* mutant indicating not only the amino acid sequence but also the function of EOL-1 is conserved (Shen et al., 2014). *eol-1* is one of 10 *C. elegans* DXO homologues. Interestingly, 6 of these genes (M01G12.7, M01G12.9, M01G12.14, Y47H10A.3, Y47H10A.4, and Y47H10A.5) are clustered as tandemly duplicated genes, adjacent to *rrf-2*, one of four *C. elegans* Rdrps, and one (C37H5.14) is adjacent to *hsp-6*.

To verify the transcriptional upregulation of *eol-1* in *hsp-6(mg585)* mutant and test if the induction requires DRH-1, a transcriptional fusion reporter *Peol-1::GFP* containing the *eol-1* promoter, GFP and *eol-1* 3’UTR was constructed. In wild type animals treated with control RNAi, *Peol-1::GFP* was barely detectable (Figure 2D and 2E); in the *hsp-6(mg585)* mitochondrial mutant, *Peol-1::GFP* was strongly induced (Figure 2D and 2E). The induction of *Peol-1::GFP* was abrogated in *hsp-6(mg585)* treated with *drh-1* RNAi (Figure 2D and 2E). Since the *Peol-1::GFP* transcriptional fusion reporter is a transgene that is subject to enhanced RNAi transgene silencing by *hsp-6(mg585)* and suppressed by *drh-1*, it might not reflect the accurate expression level of *eol-1* in these mutants. To evaluate the actual mRNA level of chromosomal *eol-1*, quantitative reverse transcription PCR (RT-qPCR) was performed. The mRNA level of chromosomal *eol-1* was 30 fold increased in the *hsp-6(mg585)* single mutant, and this induction was almost completely abolished in the *hsp-6(mg585); drh-1(tm1329)* double mutant (Figure 2F). Thus, mitochondrial dysfunction of *hsp-6(mg585)* triggers the transcriptional activation of *eol-1* in a DRH-1-dependent manner.

In *C. elegans*, the mitochondrial unfolded protein response (UPR^mt^) is a transcriptional program responding to mitochondrial dysfunction and essential for mitochondrial recovery, immunity, detoxification and aging (Lin and Haynes, 2016). RNAi of the genes *atfs-1* or *dev-1*, which encode transcription factors that mediate the expression of UPR^mt^ genes (Nargund et al., 2012; Tian et al., 2016), suppress the induction of *Peol-1::GFP* (Figure 2D and 2E). We then tested if DRH-1 contributes to the activation of UPR^mt^ by observing the induction of *Phsp-6::GFP*, the canonical reporter of UPR^mt^. The *Phsp-6::GFP* was strongly induced in the *hsp-6(mg585)* mutant, and suppressed by RNAi of *atfs-1* or *dev-1* as expected (Figure 2G and 2H). RNAi of *drh-1* in the *hsp-6(mg585); Phsp-6::GFP* strain did not disrupt induction of *Phsp-6::GFP*; in fact, induction of *Phsp-6::GFP* by *hsp-6(mg585)* after *drh-1(RNAi)* was more than in *hsp-6(mg585)* alone, perhaps because *drh-1* inactivation inhibited *hsp-6(mg585)*-induced enhanced RNAi and transgene silencing. Thus, the transcriptional activation of *eol-1* requires UPR^mt^ signaling, whereas DRH-1 is not part of the general UPR^mt^.

The DRH-1 dependent upregulation of *eol-1* implies that EOL-1 might act in the enhanced RNA interference pathway induction of the *hsp-6(mg585)* mutant. To test this possibility, a loss-of-function mutant *eol-1(mg698)* was generated by CRISPR-Cas9 (Arribere et al., 2014). In *eol-1(mg698)*, six nucleotides “TGATCA”, which contains a stop codon and a Bcl I endonuclease recognition sequence to facilitate genotyping, was inserted into the *eol-1* locus and resulted in a “Lys25 to stop” nonsense mutation (Figure 3A). The *rol-6(su1006)* and *sur-5::GFP* transgenes were tested for silencing in the *hsp-6; eol-1* double mutant. For the *rol-6(su1006)* transgene, the *hsp-6(mg585); eol-1(mg698)* double mutant showed 82% rolling compared to 27% in the *hsp-6(mg585)* single mutant (Figure 3B). For the *sur-5::GFP* transgene, the GFP signal in the *hsp-6(mg585); eol-1(mg698)* double mutant was substantially increased relative to the *hsp-6(mg585)* single mutant (Figure 3C and 3D). Moreover, the Eri phenotype of *hsp-6(mg585)* as assessed using the lethality of the *lir-1* or *his-44* dsRNAs was also suppressed by *eol-1(mg698)* (Figure 3E).

**Figure 3.**
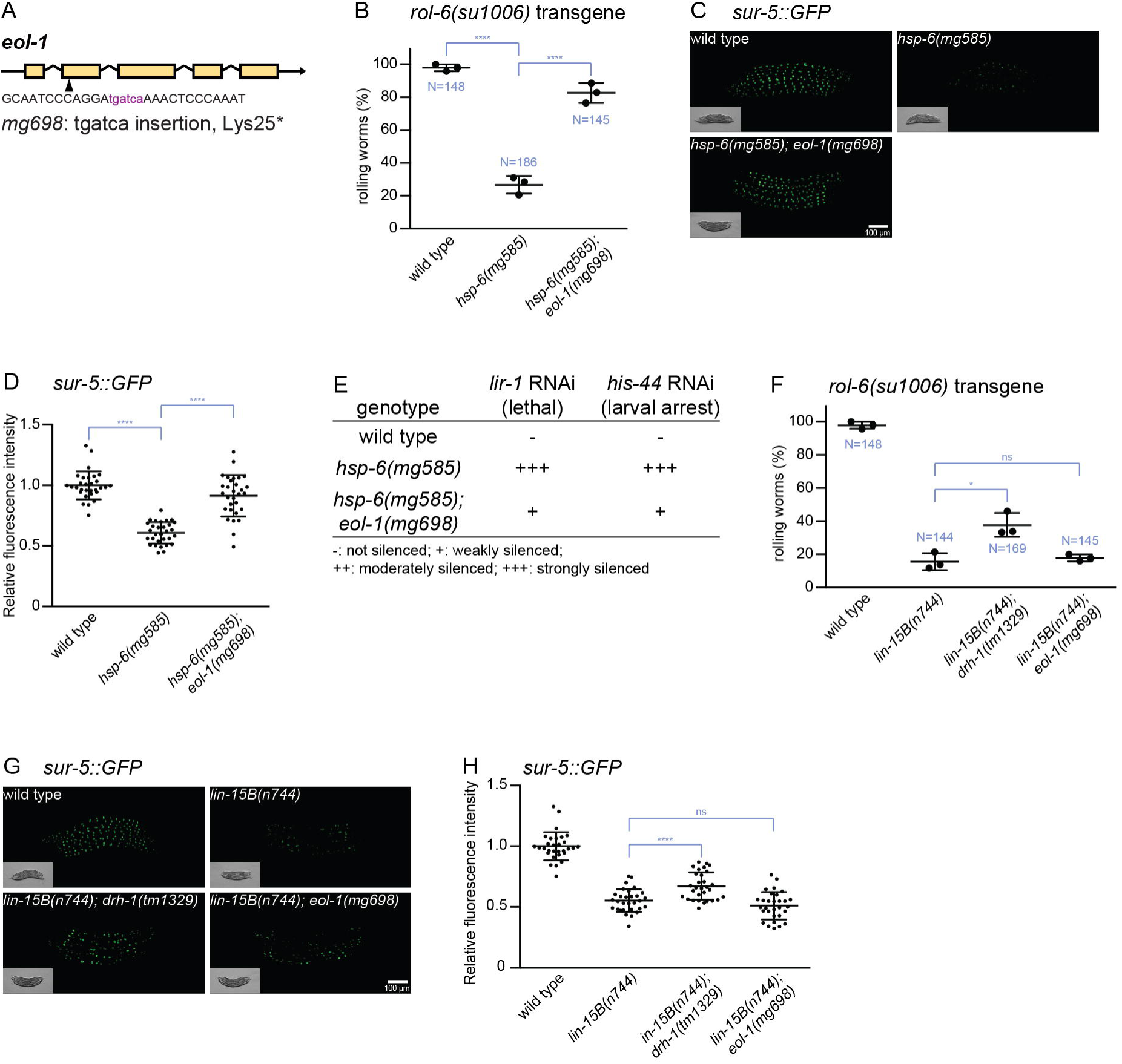
EOL-1 gene activity is required for enhanced RNAi. (A) The *eol-1(mg698)* mutant allele generated by CRISPR-Cas9. (B) Transgene silencing test with *rol-6(su1006)* transgene. The expression of the *rol-6* collagen mutation from the transgene is silenced by enhanced RNAi in *hsp-6(mg585)* mutant, and this transgene silencing depends on *eol-1* gene activity. Results of 3 replicate experiments are shown. N: total number; ****p<0.0001. (C) and (D) Transgene silencing test with *sur-5::GFP* transgene. This transgene was ubiquitously expressed in all somatic cells. The expression of the *sur-5::GFP* from the transgene is silenced by enhanced RNAi in the *hsp-6(mg585)* mutant, and this transgene silencing requires *eol-1*. Animals were imaged in (C) and the fluorescence was quantified in (D). ****p<0.0001. (E) Enhanced RNAi response to *lir-1* RNAi or *his-44* RNAi in *hsp-6(mg585)* mutant requires *eol-1*. RNAi of *lir-1* or *his-44* causes lethality/arrest on *hsp-6(mg585)* mutant but not wild type. The enhanced RNAi is suppressed by the *eol-1(mg698)* mutation. (F) Silencing of *rol-6(su1006)* transgene caused by synMuvB enhanced RNAi mutations does not depend on *eol-1*. The *rol-6(su1006)* transgene is silenced by the synMuvB *lin-15(n744)* mutation, and this transgene silencing was slightly suppressed by *drh-1(tm1329)*, but not by *eol-1(mg698)*. Results of 3 replicate experiments are shown. N: total number; *p<0.05; ns p>0.05. (G) and (H) Silencing of *sur-5::GFP* transgene caused by synMuvB enhanced RNAi mutations does not require *eol-1*. The *sur-5::GFP* transgene is silenced by the synMuvB *lin-15(n744)* mutation, and this transgene silencing was slightly suppressed by *drh-1(tm1329)*, but not *eol-1(mg698)*. Animals were imaged in (G) and the fluorescence was quantified in (H). ****p<0.0001; ns p>0.05.

Because mammalian DXO, a homologue of *C. elegans* EOL-1, is involved in mRNA decay, we tested whether EOL-1 mediates the degradation of mRNAs from transgenes. In such a model, *eol-1* might also be required for the somatic transgene silencing caused by other Eri mutants, for example, by the large synMuv B class of Eri mutants. Although *drh-1(tm1329)* slightly suppressed the transgene silencing of a *lin-15B* null mutant, *eol-1* did not contribute to transgene silencing by the synMuvB mutants (Figure 3F, 3G and 3H). Thus, EOL-1 does not act in the synMuvB RNAi pathway that silences transgenes and is specifically required for mitochondrial dysfunction-induced transgene silencing.

### Formation of EOL-1 foci requires biogenesis of mitochondrial RNAs

In order to understand the function of DRH-1 and EOL-1 in silencing of transgenes, we monitored their subcellular localization by fusing mScarlet at the N-terminus of DRH-1 and C-terminus of EOL-1, respectively, under the control of *rpl-28* promoter in a miniMOS vector (Frokjaer-Jensen et al., 2014). The miniMOS generated single copy transgene is able to avoid multicopy transgene silencing by the RNAi pathway, and the ribosome promoter *Prpl-28* is a constitutive promoter with universal expression in all tissues. In wild type animals, mScarlet::DRH-1 and EOL-1::mScarlet were localized diffusely in the cytosol without any notable pattern (Figure 5A). In the *hsp-6(mg585)* mitochondrial mutant, strikingly, EOL-1::mScarlet formed massive puncta in many cell types (Figure 5A). mScarlet::DRH-1 remained diffusely localized in *hsp-6(mg585)* (Figure 5A), just as no change was observed in DRH-1::GFP subcellular distribution upon Orsay virus infection (Ashe et al., 2013). The formation of EOL-1::mScarlet foci in the *hsp-6* mitochondrial mutant, and the specificity of EOL-1 in the suppression of *hsp-6(mg585)* induced silencing, and the mammalian finding that MDA5 recognizes mitochondrial dsRNAs generated a hypothesis that the target of EOL-1 decapping are RNAs derived from the mitochondrial genome. First, we examined whether the cytoplasmic EOL-1 foci localized at the mitochondria surface. Because loss-of-function *eol-1(mg698)* suppresses the mitochondrial defect-induced *rol-6(su1006)* transgene silencing of in the hypodermis, the hypodermal-specific *col-10* promoter driven TOMM-20::GFP miniMOS single copy transgene was used to monitor mitochondria in hypodermal cells. However, EOL-1::mScarlet foci were not specifically associated with the mitochondria (Figure 5B). Therefore, EOL-1 accumulates as foci in response to mitochondrial dysfunction, but these foci as not mitochondrially-associated and may function in the cytosol.

**Figure 4.**
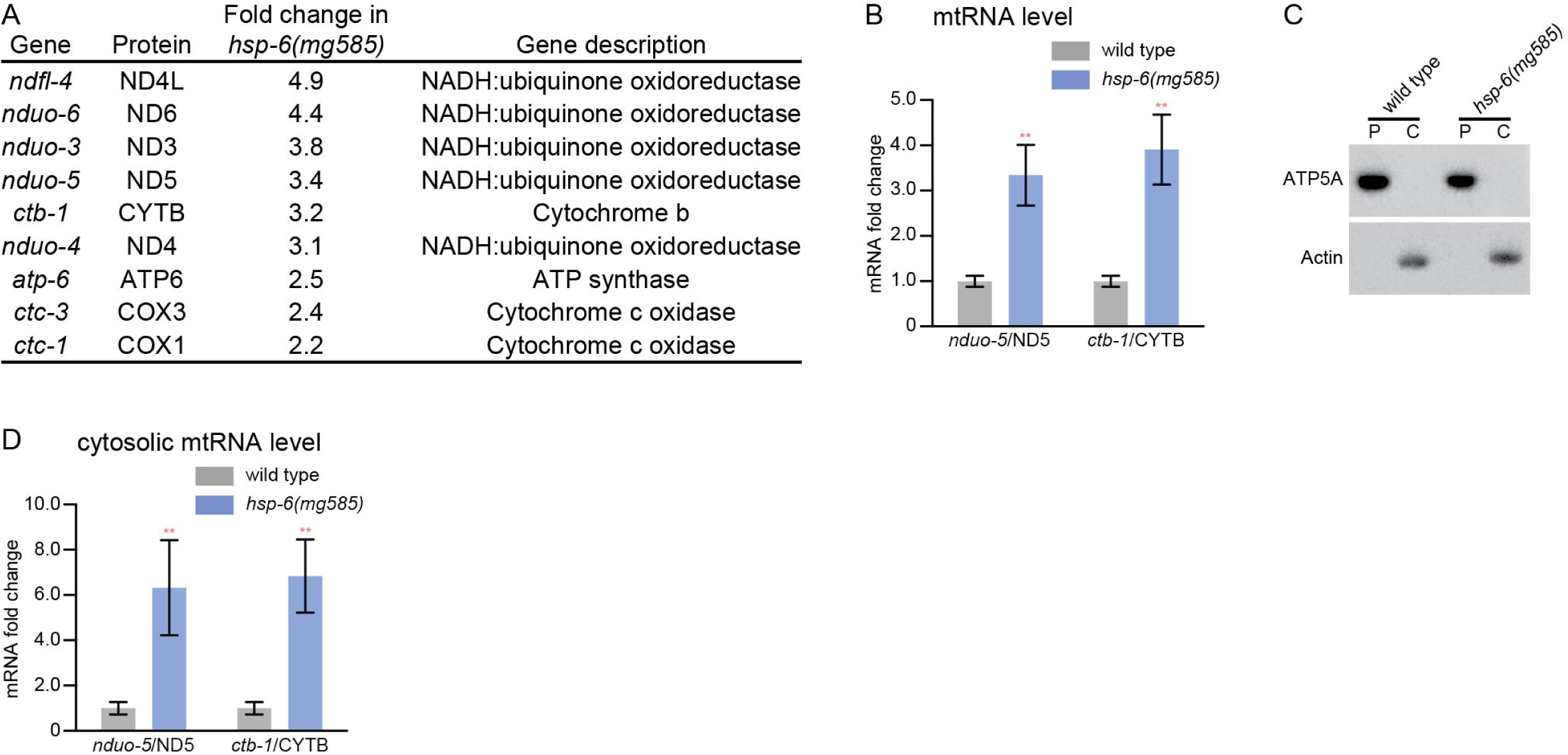
Release of mitochondrial RNA into the cytosol. (A) Electron transport chain genes encoded in the mitochondrial genome are induced by the *hsp-6(mg585)* mitochondrial chaperone gene mutation. Of the 12 genes in the mitochondrial genome that encode subunits of electron transport chain, 9 genes were upregulated in *hsp-6(mg585)* mutation. (B) The mRNA level of *nduo-5*/ND5 or *ctb-1*/CYTB was induced by *hsp-6(mg585)* mitochondrial mutation. Both *nduo-5*/ND5 (NADH:ubiquinone oxidoreductase) and *ctb-1*/CYTB (cytochrome b) are transcribed from mitochondrial genome and their expression level was evaluated by RT-qPCR assays. **p<0.01. (C) and (D) The mRNA level of *nduo-5*/ND5 or *ctb-1*/CYTB in the cytosol was increased in *hsp-6(mg585)* mitochondrial mutation. The pellet and cytosolic fractions were separated by centrifugation and probed with ATP5A and actin antibodies, respectively to assess purification in (C). The mRNA level of *nduo-5*/ND5 or *ctb-1*/CYTB in the mitochondria free cytosolic fraction was evaluated by RT-qPCR in (D). P: pellet fraction; C: cytosolic fraction.

**Figure 5.**
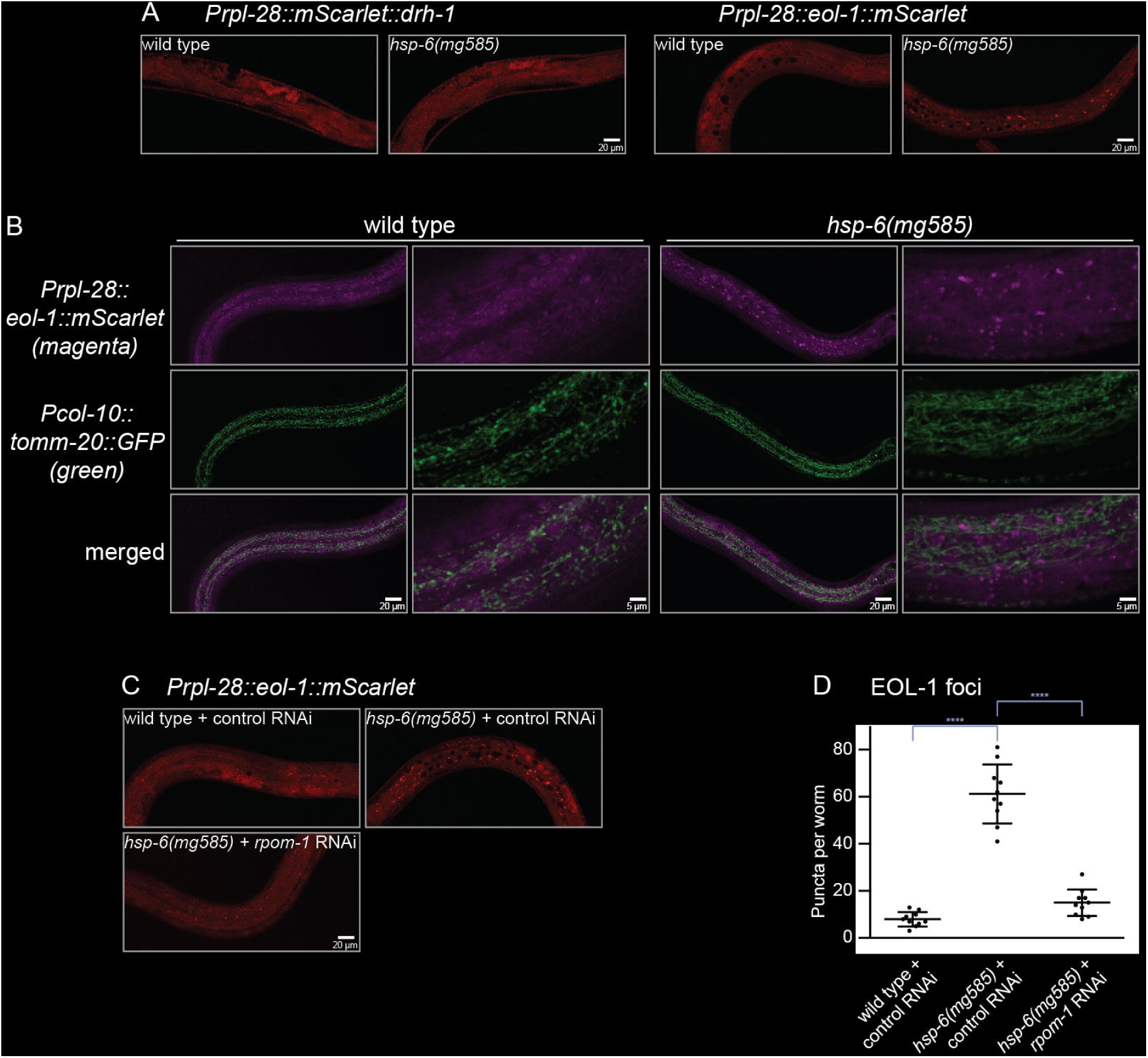
EOL-1 protein forms cytoplasmic foci if mitochondria are dysfunctional. (A) EOL-1 protein, but not DRH-1 protein form easily seen puncta in the *hsp-6(mg585)* mitochondrial mutant but not in wild type. Mitochondrial dysfunction caused by *hsp-6(mg585)* mutation trigged the formation of EOL-1::mScarlet foci, while mScarlet::DRH-1 remained diffusely localized. (B) EOL-1 foci are not associated with the mitochondria. In the *hsp-6(mg585)* mutant, the EOL-1::mScarlet foci do not colocalize with mitochondria that were visualized by the mitochondrial outer membrane protein TOMM-20::GFP. (C) and (D) Formation of EOL-1 cytoplasmic foci requires the transcription of mitochondrial RNA from mitochondrial genome by RPOM-1 RNA polymerase. The EOL-1::mScarlet foci formed in the *hsp-6(mg585)* mutant were inhibited by RNAi of mitochondrial RNA polymerase *rpom-1*/ POLRMT. Animals were imaged in (C), and the number of foci was quantified in (D). ****p<0.0001.

We explored whether *C. elegans* mitochondria release RNA into the cytosol, as has been observed in the MDA5 mitochondrial surveillance pathway in human and *D. melanogaster* (Dhir et al., 2018; Pajak et al., 2019). The human mtDNA transcribes 37 genes, including 13 mRNAs encoding subunits of electron transport chain (ETC), 2 ribosomal RNAs (rRNA) and 22 transfer RNAs (tRNA), residing on both the heavy and light strands of the mitochondrial DNA (Hallberg and Larsson, 2014). Transcription on both strands generated two genome size polycistronic transcripts that are processed into individual genes. During this processing, the complementary noncoding RNAs in mammals are degraded by the mitochondrial degradosome formed by RNA helicase SUV3 and polynucleotide phosphorylase PNPase (Borowski et al., 2013). Loss of SUV3 or PNPase stabilize the noncoding RNAs to form dsRNAs with the genes derived form the opposite strand (Dhir et al., 2018). The *C. elegans* mitochondrial genome encodes 36 genes with one less ETC subunit (Okimoto et al., 1992). However, these 36 genes are encoded exclusively on the heavy strand of *C. elegans* mtDNA and no transcription of the light strand has been detected (Blumberg et al., 2017), which disfavors the mitochondrial dsRNA model.

mRNA-seq analysis of *hsp-6(mg585)* revealed that 9 of 12 mitochondrial mRNA encoding ETC subunits were upregulated (Figure 4A). The increased level of mitochondrial mRNA was further verified by RT-qPCR analysis of *nduo-5*/ND5 and *ctb-1*/CYTB (Figure 4B). In order to examine if the increased abundance of mitochondrial mRNAs within the mitochondrial matrix caused their release into the cytosol, the cytosolic fraction (actin antibody was used as a control protein for this fractionation) was separated by centrifugation from the pellet fraction containing mitochondria (ATP5A antibody used as a control for this fractionation) (Figure 4C). RT-qPCR analyses showed that the mRNA levels of *nduo-5*/ND5 and *ctb-1*/CYTB were substantially increased in the cytosol of *hsp-6(mg585)* compared to wild type (Figure 4D).

The strategy of single strand transcription from the mitochondrial genome is not just parochial to *C. elegans*; it is a strategy used by many members of the genus *Caenorhabditis* (Li et al., 2018). If the unique strand transcriptional architecture of the *Caenorhabditis* mitochondrial genome is linked with production of dsRNA from the mitochondria, and if the release of mitochondrial RNA into the cytosol in mitochondrial mutants is the trigger for the formation of EOL-1 foci, we would expect a suppression of EOL-1 focus formation by shutting down mitochondrial RNA biogenesis. The human gene POLRMT encodes a mitochondrial DNA-directed RNA polymerase that catalyzes the transcription of mitochondrial DNA (Hallberg and Larsson, 2014). RNAi of *rpom-1*, the *C. elegans* orthologue of human POLRMT, potently reduced the number of EOL-1 foci in *hsp-6(mg585)* mutant (Figure 5C and 5D). Therefore, the formation EOL-1 foci required the biogenesis of mitochondrial RNA. Mitochondrial RNA polymerase is central to all gene expression in the mitochondrion, including the 12-13 protein coding genes, the tRNA genes and the rRNA genes. So a defect in *rpom-1* is expected to be pleiotropic and thus difficult to ascribe to production of dsRNA from mitochondrial transcripts. But paradoxically, with 36 client genes in the mitochondrion, *rprom-1* has a rather circumscribed sphere of genes it controls, so the its function upstream of EOL-1 foci could very well be via dsRNAs produced from mitochondria but only released in mitochondrial mutant strains.

### Enhanced RNA interference mediates lifespan extension in mitochondrial mutant strains

While the majority of mitochondrial components are essential for eukaryotes and null alleles of most mitochondrial genes are lethal, *C. elegans* reduction-of-function mitochondrial mutants such as *hsp-6(mg585)* (Figure 6A and 6D) increase longevity, in some cases dramatically (Dillin et al., 2002; Lee et al., 2003). We explored whether the enhanced RNAi output of the *hsp-6(mg585)* mitochondrial mutant contributes to its longevity. *drh-1(tm1329)* or *eol-1(mg698)* single mutants did not shift the survival curve compared to wild type (Figure 6B and 6D), showing that these mutants are not short-lived or sickly in some way. But *drh-1(tm1329)* or *eol-1(mg698)* mutations strongly suppressed the lifespan extension of *hsp-6(mg585)*, showing that *drh-1* or *eol-1* gene activities are required for the extension of lifespan by *hsp-6(mg585)* (Figure 6C and 6D). Because *drh-1* and *eol-1* mediate the enhanced RNA interference, an antiviral response, of mitochondrial mutants, and because these outputs also mediate in the increased longevity of mitochondrial mutants, but not the normal longevity of wild type, these data support the model that enhanced antiviral defense as a key anti-aging output from mitochondrial mutations.

**Figure 6.**
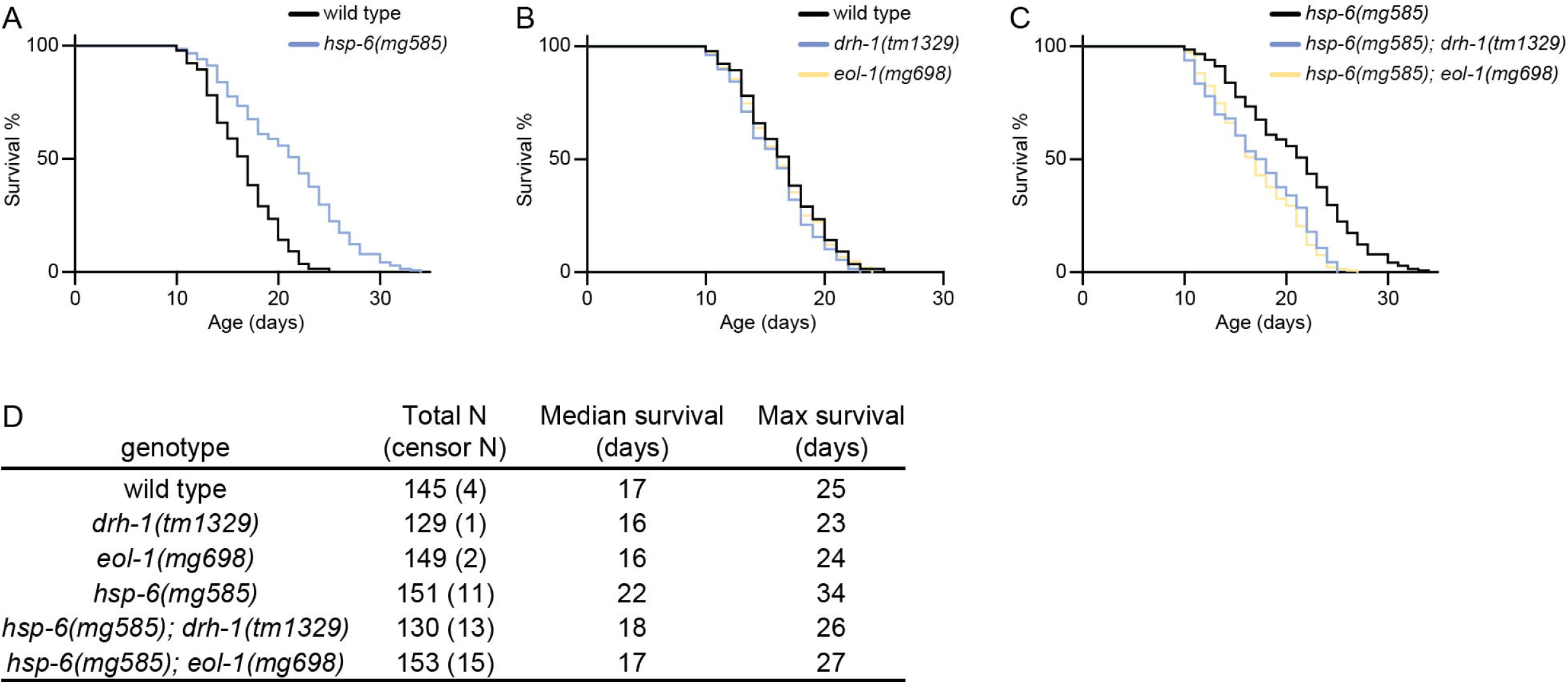
Anti-aging effect of RNAi pathway. (A) The *hsp-6(mg585)* mitochondrial mutation extends lifespan. (B) *drh-1(tm1329)* or *eol-1(mg698)* single mutants do not extend lifespan. (C) *drh-1(tm1329)* or *eol-1(mg698)* suppresses the extension of lifespan caused by *hsp-6(mg585)* mutation. (D) Median and maximal survival and number of animals surveyed and censored due to losses during the month long assay.

## Discussion

The RNAi pathway drives gene silencing of mRNAs that have signatures of being foreign. For example, many of the enhanced RNAi mutants mediate the silencing of integrated viruses in the *C. elegans* genome that are recently acquired (Fischer and Ruvkun, 2020). These viral genes are recognized as foreign by their small number of introns, for example, and by poor splice sites (Newman et al., 2018).

The antiviral defense of *C. elegans* is more closely related to that of fungi and plants than most animals, for example Drosophila and vertebrates. Nematodes, plants, and most fungi use the common pathway genes Dicer, specialized Argonaute/PIWI proteins, and most distinctively, RNA dependent RNA polymerases to produce primary siRNAs and secondary siRNAs against invading viruses. Because these pathways are common across many (but not all) eukaryotes, it is likely that the common ancestor of animals, plants, fungi, and some protists carried the Dicer, Argonaute, and Rdrp genes that mediate RNA interference defense. RNA dependent RNA polymerase genes, paradoxically, are almost always carried by RNA viruses to mediate their replication (Smith et al., 2014). Thus a key protein in antiviral defense in most eukaryotes, is a protein that may spread across eukaryotes from viral infections.

The loss of Rdrp genes in most animals species appears to be associated with the evolution of the Sting and NFkappaB interferon RNA virus defense pathway of most insects and vertebrates but not in nematodes or plants or fungi. In addition to the interferon pathway, the piRNA pathway has specialized in insects and vertebrates to mediate antiviral defense (Aravin et al., 2007). The piRNA pathway is not present in fungi or plants and appears to not have become as central to viral defense in *C. elegans* which has not lost its RNA dependent RNA polymerase, like most other animal species such as vertebrates and insects (Tabach et al., 2013). Viral defense in *C. elegans* is strongly induced in the many mutants that cause enhanced RNA interference, *eri-1* to *eri-10*, the many synMuvB mutations, and the mitochondrial mutants we characterize here. These mutations may induce antiviral defense because viral infections may compromise these genetic pathways or because these mutations genetically trigger an antiviral state that is normally under physiological regulation. A surveillance system to monitor the states of these pathways may be an early detection system for viruses to induce expression and activities of Dicer, Argonaute proteins and RNA dependent RNA polymerases in siRNA production, targeting, and amplification (Govindan et al., 2015; Melo and Ruvkun, 2012).

Regardless of whether antiviral defense is mediated by Rdrps in nematodes or by MAVS and interferon signaling, in vertebrates and most insects, the mitochondria are woven into the pathways. For Flock House Virus and other nodaviruses, the mitochondrion is a center of viral replication; mutations that compromise the mitochondrion in yeast for example, cause defects in antiviral responses (Hao et al., 2014; Miller and Ahlquist, 2002; Miller et al., 2001).

It is possible that viruses generally need the oxidizing power of O_2_ for the massive production of disulfides in virion assembly in the ER. Thus mitochondrial dysfunction may be a key pathogenic feature of viral infection and systems have evolved to trigger antiviral defense if mitochondria dysfunction is detected. The mitochondria has also been implicated in the Drosophila and mammalian antiviral piRNA pathway: in Drosophila, retroviruses are recognized by the dedicated piRNA pathway that generates piRNAs from integrated flamenco as well as other retroviral elements (Malone et al., 2009) to target newly encountered viruses that are related to these integrated viruses. A variety of mitochondrial outer membrane proteins couple piRNAs to target loci (Huang et al., 2011; Munafo et al., 2019). And the Sting interferon pathway also uses the MAVS and RIG-I interaction with mitochondria to signal antiviral responses.

We have shown that enhanced RNA interference is also induced by many mutations that compromise mitochondrial function. Human mitochondrial mutations also trigger antiviral immunity (Dhir et al., 2018; West et al., 2015). The human RIG-I RNA helicase recognizes viral dsRNA or ssRNA 5’-triphosphate (Chow et al., 2018). Our data suggests that the signal that triggers the homologous RNA helicase, DRH-1, are mitochondrial dsRNAs that are released to the cytosol. The EOL-1/DXO exonuclease may remove the 5’ cap of mitochondrial RNA, such as a 5’-NAD modification (Bird et al., 2018), to facilitate the recognition by DRH-1. The 5’ ends of the majority of messenger RNAs (mRNA) are chemically modified or capped to protect from RNA nucleases (Grudzien-Nogalska and Kiledjian, 2017). Prokaryotic and eukaryotic cells have different strategies to modify their mRNA with di- or triphosphate in bacteria and N^7^-methyl guanosine (m^7^G) cap in eukaryotes. Thus the eukaryotic 5’ cap is a conspicuous sign to identify self, rather than the non-self uncapped RNA from virus or bacteria.

A decrement in mitochondrial function is one of the most potent mechanisms to increase longevity in a variety of species (Lee et al., 2003). Our analysis of the *eol-1* and *drh-1* pathway from mitochondrial dysfunction to enhanced RNA interference and antiviral activity is a key output from mitochondria for anti-aging. This would suggest that the dramatic increase in frailty in old age could reflect viral vulnerability. In fact, one of the most dramatic hallmarks of the recent Covid-19 epidemic is that the virus is 1000x more lethal to the elderly than young adults. In support of this model, in mammals, a reduction in Dicer expression in heart, adipose and brain is a hallmark of aging (Kaneko et al., 2011; Tarallo et al., 2012). A loss-of-function mutant of DCR-1/Dicer shortens *C. elegans* lifespan (Mori et al., 2012).

Anti-bacterial immune responses are also activated upon mitochondrial disruption (Liu et al., 2014; Mao et al., 2019; Pellegrino et al., 2014), suggesting that perhaps the *C. elegans* antiviral and antibacterial responses may have common features. *eol-1* was discovered based on its effect on olfactory learning upon *P. aeruginosa* infection (Shen et al., 2014), and it is induced by particular bacterial pathogens (Sowa et al., 2020). Gene expression profiles suggest that after virus infection, loss of *drh-1* causes a shift from antiviral response to antibacterial response (Sarkies et al., 2013). Thus, the immune network in *C. elegans* may be coupled to the surveillance of many cellular systems to trigger a variety of defense responses, including enhanced RNA interference. The enhanced RNAi mutants may constitutively express these normally transient states of antiviral or antibacterial defense.

## Materials and Methods

### *C. elegans* maintenance and strains used in this study

All C. elegans strains were cultured at 20 °C. Strains used in this study are listed in Supplemental Table S2.

### Generation of transgenic animals

For *Peol-1::GFP*, the plasmid harboring [*Peol-1::GFP::eol-1 3’UTR + cbr-unc-119(+)*] was injected into *unc-119(ed3)*. For single copy transgenes *Prpl-28::mScarlet::drh-1, Prpl-28::eol-1::mScarlet* and *Pcol-10::tomm-20::GFP*, the plasmid was injected into *unc-119(ed3)* following the miniMOS protocol (Frokjaer-Jensen et al., 2014). For CRISPR of *eol-1(mg698)*, we chose *dpy-10(cn64)* as the co-CRISPR marker (Arribere et al., 2014), and pJW1285 (Addgene) to express both of guide-RNA (gRNA) and Cas9 enzyme (Ward, 2015).

### Microscopy

The fluorescent signals of *sur-5::GFP, Peol-1::GFP* and *Phsp-6::GFP* transgenic animals were photographed by Zeiss AX10 Zoom.V16 microscope. The subcellular pattern of *Prpl-28::mScarlet::drh-1, Prpl-28::eol-1::mScarlet* and *Pcol-10::tomm-20::GFP* were carried out on the Leica TCS SP8 confocal microscope. Photographs were analyzed by Fiji-ImageJ.

### Sample collection and RNA isolation

For estimation of the mRNA level of *eol-1, nduo-5* and *ctb-1*, around 200 L4 larvae were hand picked from mixed population and frozen by liquid nitrogen. Total RNA was isolated by TRIzol extraction (Thermo Fisher #15596026). For analyses of cytosolic mRNA level of *nduo-5* and *ctb-1*, approximately 2000 worms were synchronized by bleach preparation of eggs, hatching progeny eggs to L1 larvae, grown till L4 stage and frozen by liquid nitrogen. Worm lysates were generated by TissueLyser with steel beads (Qiagen #69989), and re-suspended in 500 μl of the lysis buffer containing 0.8 M sucrose, 10mM Tris-HCl, 1 mM EDTA, 1 X protease inhibitor cocktail (Roche 11873580001) and 1 U/μl murine RNase inhibitor (New England BioLabs, #M0314L). The lysate was centrifuged at 2500g for 10 min at 4 °C to remove the pellet debris. And the supernatant was centrifuged at 20000g for 10 min at 4 °C. Collect the pellet for western blot and the supernatant was centrifuged at 20000g for 10 min at 4 °C. Take 100 μl of the supernatant for western blot, and 300 μl for RNA isolation with TRIzol.

### RT-qPCR

The cDNA was generated by ProtoScript^®^ II First Strand cDNA Synthesis Kit (New England BioLabs, #E6560L). qPCR was performed toward *eol-1, nduo-5* and *ctb-1* with *act-1* as control by iQ™ SYBR^®^ Green Supermix (BIO-RAD, #1708880).

### Western blot

The pellet or supernatant from the previous centrifugation were mix NuPAGE™ LDS sample buffer (Thermo Fisher #NP0007) and heated at 70 °C for 10 minutes. Samples were loaded onto NuPAGE™ 4-12% Bis-Tris Protein Gels (Thermo Fisher #NP0323BOX) and run with NuPAGE™ MES SDS running buffer (Thermo Fisher #NP0002). After semi-dry transfer, PVDF membrane (Millipore #IPVH00010) was blocked with 5% nonfat milk, and probed with anti-ATP5A (Abcam ab14748) or anti-actin (Abcam #ab3280) primary antibodies. The membrane was developed with SuperSignal™ West Femto Maximum Sensitivity Substrate (Thermo Fisher #34096) and visualized by Amersham Hyperfilm (GE Healthcare #28906845).

### Lifespan analysis

Animals were synchronized by egg-laying and grown until the L4 stage as day 0. Adults were separated from their progeny by manually transfer to new plates. Survival was examined on a daily basis and the survival curve was generated by GraphPad Prism.

## Author contributions

G.R. supervised the study. K.M. and G.R. designed the experiments and wrote the manuscript. K.M. performed most experiments and analyzed results. P.B. performed microinjection.

## Acknowledgements

We thank Caenorhabditis Genetics Center and National BioRescource Project (Tokyo, Japan) for providing strains. K.M. is a Damon Runyon Fellow supported by the Damon Runyon Cancer Research Foundation (DRG-2213-15). This work is supported by a grant from the National Institute of Health awarded to G.R. (NIH GM044619 and GM16636).

## Figure Legends

**Supplemental Figure 1.**
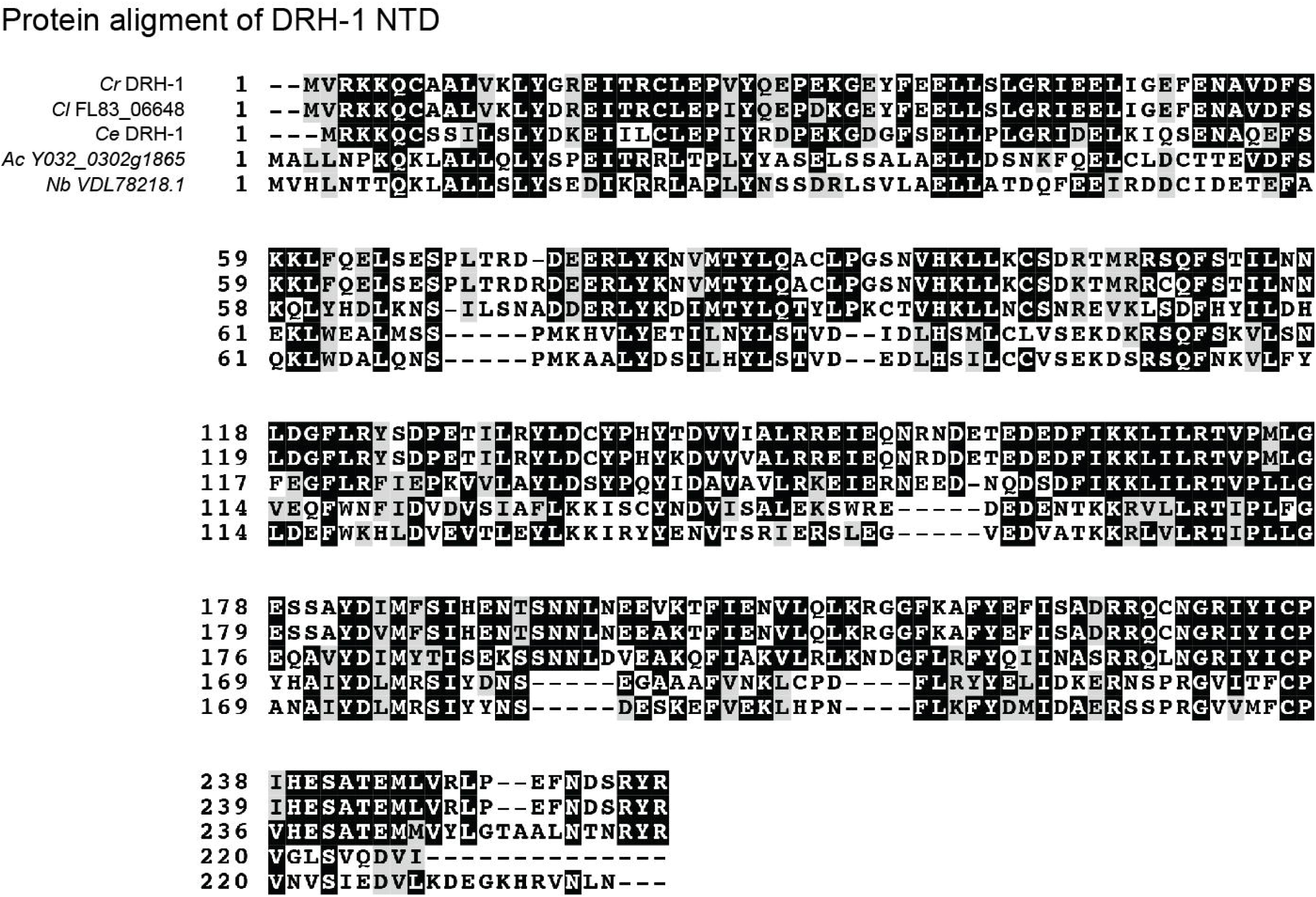
Protein alignment of DRH-1 NTD in nematode species. *Cr*: *Caenorhabditis remanei*; *Cl*: *Caenorhabditis latens*; *Ce*: *Caenorhabditis elegans*; *Ac*: *Ancylostoma ceylanicum*; *Nb*: *Nippostrongylus brasiliensis*.

